# Trading water for carbon: Sustained photosynthesis at the cost of increased water loss during high temperatures in a temperate forest

**DOI:** 10.1101/760249

**Authors:** Anne Griebel, Lauren T. Bennett, Daniel Metzen, Elise Pendall, Patrick N.J. Lane, Stefan K. Arndt

## Abstract

Forest carbon and water fluxes are often assumed to be coupled as a result of stomatal regulation during dry conditions. However, recent observations have indicated increased transpiration rates during isolated heat waves across a range of eucalypt species under experimental and natural conditions, with inconsistent effects on photosynthesis (ranging from an increase to a near total decline). To improve the empirical basis for understanding carbon and water fluxes in forests under hotter and drier climates, we measured the water use of dominant trees, and the ecosystem-scale carbon and water exchange in a mature temperate eucalypt forest over three summer seasons. The forest maintained photosynthesis within 16% of peak photosynthesis rates during all conditions, despite up to 70% reductions in canopy conductance during a 5-day heatwave. While carbon and water fluxes both decreased by 16% on exceptionally dry summer days, GPP was sustained at the cost of up to 74% increased water loss on the hottest days and during the heatwave. This led to ∼40% variation in ecosystem water use efficiency over the three summers, and ∼two-fold differences depending on the way water use efficiency is calculated. Furthermore, the forest became a net source of carbon following a 137% increase in ecosystem respiration during the heat wave, highlighting that the potential for temperate eucalypt forests to remain net carbon sinks under future climates will depend not only on their potential to maintain photosynthesis during higher temperatures, but also on responses of ecosystem respiration to changes in climate.

**Key Points:**

- GPP of temperate eucalypts was sustained at the cost of increased water use during hot periods, but both fluxes decreased during dry periods.
- WUE estimates for the same period differed up to two-fold depending on the way it was calculated.
- Doubling of ecosystem respiration turned the forest from a net sink into a net source of carbon during a longer heatwave.

## 1 Introduction

A hotter and drier future is likely for many of Australia’s ecosystems. Australia’s mean annual temperature has increased by 1 °C since 1910, temperature distributions have shifted towards higher average monthly maximum and minimum temperatures, and the duration, frequency and intensity of extreme heat events has increased (BOM 2016a). The years 2013-2015 were among the top 10 hottest years on record, including a number of significant heatwaves in southeast Australia (BOM 2013, 2014, 2015). In addition, southeastern Australia has become drier due to severe rainfall deficiencies since the year 2000 (BOM 2016b). This indicates increased potential for climate-induced stress in Australian ecosystems, given projections of warmer and drier conditions over much of the Australian continent in coming decades (IPCC 2013). This will likely result in more hot days and fewer cool days, in addition to more time spent in drought as winter and spring rainfall is predicted to decrease further (BOM 2016a).

Over 900 eucalypt species occur in a broad range of climates in Australia, some with relatively narrow distributions, which could make them vulnerable to a changing climate (Brouwers et al., 2013; Hughes et al., 1996; Jurskis, 2005), and especially to extreme climate events (Choat et al., 2012; Matusick et al., 2013; Mitchell et al., 2014a). Many eucalypt species close their stomata to prevent excessive water loss in response to dry conditions (Breshears et al., 2013; Eamus et al., 2008), which delays embolisms in the stem xylem (Choat et al., 2012; Tyree & Sperry, 1989) at the cost of decreases in photosynthesis and increases in the vulnerability of leaves to heat and light stress (McDowell et al., 2008; McDowell, 2011; Mitchell et al., 2014b; Thomas & Eamus, 1999; Whitehead & Beadle, 2004). Stomatal conductance varies with supply and demand for CO2 by photosynthesis (intercellular CO2 concentration), leaf irradiance and leaf temperature, as well as atmospheric vapor pressure deficit and leaf turgor (Ball et al., 1987; Cowan, 1978; Medlyn et al., 2011; Tuzet et al., 2003). Hence, photosynthesis, transpiration and stomatal conductance are commonly assumed to be coupled; that is, photosynthesis and transpiration both decrease with increasing stomatal regulation under most environmental conditions (Farquhar & Sharkey, 1982; Leuning, 1995; Tuzet et al., 2003). However, isolated studies provide evidence of a decoupling of photosynthesis from stomatal conductance in some tree species during extreme heat stress (Ameye et al., 2012; De Kauwe et al., 2019; Drake et al., 2018; Urban et al., 2017). For example, photosynthesis decreased in seven eucalypt forests (De Kauwe et al., 2019) or was near zero in 1- year-old *E. parramattensis* saplings (Drake et al., 2018) as water loss increased under high temperatures. This indicates that latent cooling of leaves by transpiration might be an important mechanism to cope with extended heat stress (Drake et al., 2018), in turn affecting the plant’s carbon assimilation rate per unit stomatal conductance as well as the plant’s water use efficiency (WUE). Nonetheless, in other temperate forest types dominated by eucalypts, photosynthesis increased with transpiration during a single heat wave event (van Gorsel et al., 2016), highlighting the need for further studies of concurrent carbon and water fluxes across an extended range of weather conditions.

Individual studies of transpiration cooling under heat stress have largely focused on young plants (Ameye et al., 2012; Drake et al., 2018; Urban et al., 2017), whereas direct measures of stomatal conductance, carbon assimilation (GPP) and transpiration at the ecosystem level remain challenging. Canopy conductance (Gc, as an approximation of stomatal conductance) is commonly derived by inverting the Penman-Monteith equation using directly measured latent heat flux (LE; Monteith, 1965). WUE has been estimated as the sum of GPP over the sum of ET, but this does not account for the non-linear response of LE to vapor pressure deficit (VPD). Thus, alternative formulations of ecosystem-scale WUE as underlying WUE (Zhou et al., 2015, 2014), or as intrinsic WUE (Beer et al., 2009; Lloyd et al., 2002; Schulze and Hall, 1982) have been suggested to more accurately understand the underlying physiological mechanisms.

Studies examining concurrent carbon and water fluxes under heat stress in natural mature eucalypt forests are limited to isolated heatwave events (De Kauwe et al., 2019; van Gorsel et al., 2016). Further, it remains unclear if eucalypts respond differently to multi-day heatwaves compared with individual hot days, and how such responses might be mediated by water availability. As this is currently neither well understood, nor integrated into process-based ecosystem models, it limits the potential to predict how future climates characterized by more frequent and intense heatwaves will influence the physiology, productivity, and distribution of temperate forest eucalypts. We combined three years of concurrent sap flow and eddy covariance summer measurements in a natural temperate eucalypt forest to examine the dynamics of photosynthesis and water use during the hottest days, the driest days, and a 5-day heatwave. We hypothesized that (i) photosynthesis and transpiration would both decrease on the driest days and both increase on the hottest days (assuming no water limitations); and (ii) a longer heatwave would result in decreased carbon uptake and increased water loss, as photosynthesis decreases, and evapotranspiration increases during continuous temperature stress (assuming no water limitations). Our results have globally relevant implications for understanding the trade-offs between photosynthesis and water use from terrestrial ecosystems during exceptionally hot or dry conditions, which remain yet to be incorporated into plant hydraulic and land surface models.

## 2 Materials and Methods

### 2.1 Study site and climate

We recorded tree water use and ecosystem carbon and water fluxes in a temperate mixed-species eucalypt forest (Wombat State Forest) in southeastern Australia from January 2013 to November 2015. The study site is located near a ridge at 705 m elevation with gently sloping terrain to the southwest and northwest (< 8°; Griebel et al., 2016), approximately 100 km west of Melbourne, Australia, and is part of the TERN-SuperSite Network, TERN-OzFlux and FluxNet (AU-Wom), Sapfluxnet (AUS-WOM) and Dendroglobal network (AU-Wombat). Active forest management has been minimal since the late 1970s, with previous management practices including selective harvesting, low-intensity prescribed burning and firewood collection. The overstorey of this dry sclerophyll forest is dominated by three eucalypt species: *Eucalyptus obliqua L’Hér* (deep fibrous ‘stringybark’), *E. rubida H. Deane and Maiden* (smooth ‘gum’ bark), and *E. radiata Sieber ex DC* (short fibrous ‘peppermint’ bark; 70%, 21% and 9% of stand biomass, respectively; Griebel 2016), while the understory consists of sparse and patchy perennial grasses and the fern Austral bracken. The leaf area index (LAI, acknowledging that it includes leaves and woody biomass) was relatively stable in the first half of the study period (LAI ∼1.7 m^2^ m^−2^), and subsequently increased by ∼20% by the end of the study period (Griebel et al., 2017).

The climate is cool temperate, with typically cool and wet winters, and warm summers. The closest weather station to the study site (Ballarat Aerodrome, ∼20 km distance) recorded a long-term average annual temperature of 12.2 °C and an average annual rainfall of 690 mm (1908-2015). While the mean annual temperatures of the three study years were slightly below the long-term average (12.0 °C in 2013, 11.8 °C in 2014 and 10.8 °C in 2015), mean monthly maximum and minimum temperatures were generally greater than the World Meteorological Organization (WMO) reference period (1961-1990) in spring and summer (baseline based on Ballarat Aerodrome data; Fig. 1a,b). Likewise, probability distributions of the mean monthly minimum and mean monthly maximum temperatures indicated a high likelihood for warmer temperatures during each study year (Fig. 1c). The annual rainfall totals were within 90 mm of the long-term average of 690 mm (780 mm in 2013, 672 mm in 2014 and 679 mm in 2015), but the 2014 and 2015 monthly rainfall totals were consistently below the WMO reference totals in winter and spring (Fig. 1d).

**Figure 1.**
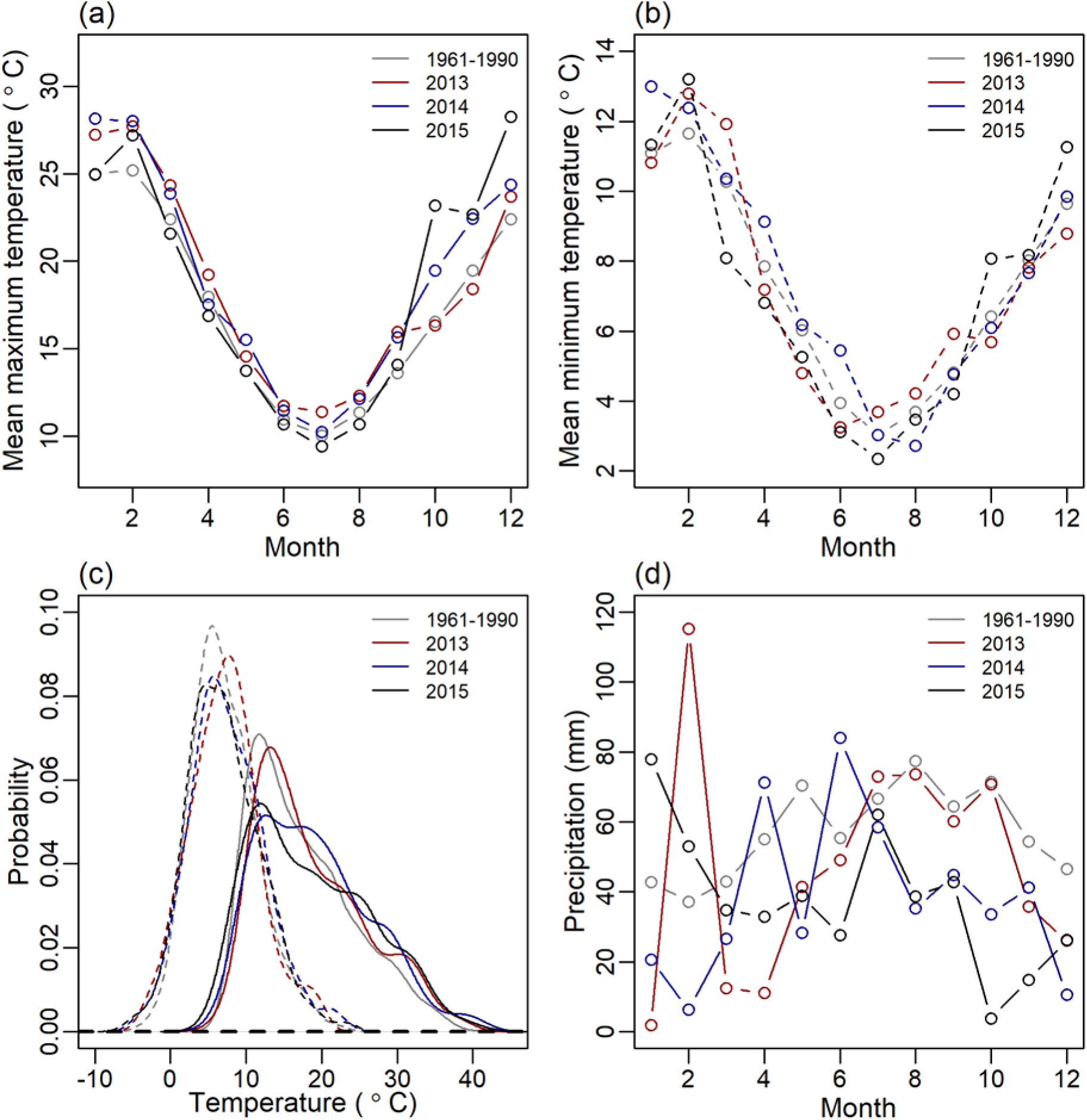
Mean monthly maximum (a) and minimum (b) temperatures in spring (months 9 to 11) and summer (months 1,2 and 12) were often higher during the three observation years than the WMO reference period (1961-1990). This affected the likelihood of higher maximum temperatures (solid lines, panel c) and to a lesser degree higher minimum temperatures (dashed lines). Rainfall distribution (monthly rainfall totals, d) during the three years was erratic during summer and autumn, but below average conditions in winter and spring in 2014 and 2015.

One of southeast Australia’s most significant heatwaves (i.e. up to 12 °C higher than the 1961-1990 January mean maximum; BOM 2014) coincided with our study period in January 2014, and involved five consecutive days reaching ∼35 °C and a peak vapor pressure deficit (VPD) of 5.4 kPa at our study site. Thus, all references to heatwaves in this paper refer to this local heatwave (‘HW’, 13 to 17 January 2014), rather than the broader-scale ‘Angry summer’ of 2013 (van Gorsel et al., 2016), which affected much of southeastern Australia but was comparatively mild at our study site due to the >700 m elevation (i.e. isolated days with maximum temperatures in the low 30s °C).

Since the 2014 heatwave was preceded by numerous rain events at the end of 2013 (i.e. was hot but well-watered), we also pooled the hottest and driest days throughout the summer months (December to February 2013-2015) for comparison with the 5-day heatwave. Here, the hottest days were those in the upper 90th percentile of maximum daily temperatures in summer (>30.7°C; 19 days), and the driest days were those in the lowest 10 percent of summer dryness as indicated by soil moisture sensors at 40 cm depth (<0.102 m^3^ m^−3^; 23 days). The 19 hottest days excluded the heatwave from 13 to 17 January 2014 and did not overlap with any of the 23 driest days. Note that soil moisture sensors at greater depths (65 cm, 1m; see ‘Climate variables and response functions’) were installed too late to capture all summer months. In addition to the WMO 1961-1990 baseline, we defined a local January baseline (‘Base’; 59 unique days) as days in January 2013, 2014 and 2015 that excluded unusual weather conditions during part of the ‘Angry summer’ from 1 to 19 January 2013, the heatwave from 13 to 17 January 2014, as well as the hottest and driest 10% of days. The incoming solar radiation was comparable between the three groups (Table 1 and Fig. S1), and the soil moisture was comparable during the baseline, the hottest days and the heatwave. Thus, apart from the driest days (which were warm and dry), the four groups primarily differed in their temperature range and associated atmospheric demand (which increased from the baseline to the driest days, the hottest days, and peaked during the heatwave; Table 1).

**Table 1.**
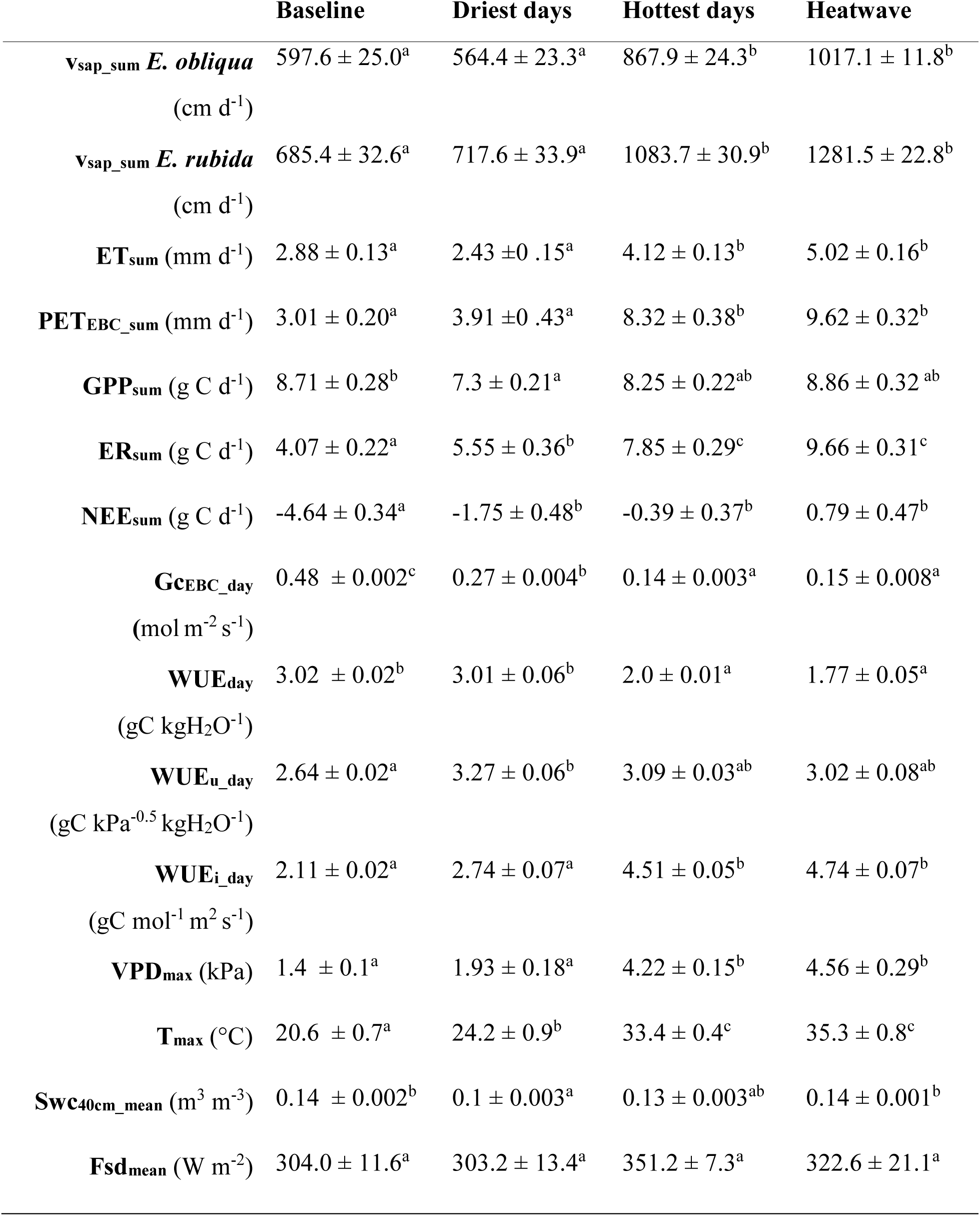
Overview of the key response and climate variables for baseline days, the hottest and the driest days, and during the heatwave. Values are mean daily sums and standard errors of water loss as sap velocity (vsap_sum) for both species and as ecosystem-scale evapotranspiration (ETsum) and potential ET (PETEBC_sum), as well as daily gross primary productivity (GPPsum), ecosystem respiration (ERsum), net ecosystem exchange (NEEsum), mean daytime canopy conductance (GcEBC_day), water use efficiency (WUEday), underlying WUE (WUEu_day = GPPxVPD^0.5^/ET) and intrinsic WUE (WUEi_day = GPP/Gc), in addition to daily maximum vapor pressure deficit (VPDmax) and temperature (Tmax), and daily mean soil water content (Swcmean) and incoming solar radiation (Fsdmean). Note that the subscript ‘EBC’ indicates that the available energy was set equal to the sum of the latent and sensible heat flux to account for the energy imbalance when inverting the Penman-Monteith combination equation to calculate canopy conductance and potential evapotranspiration. Different superscript letters indicate significant differences among the groups of days (P < 0.05, Tukey posthoc test).

### 2.2 Sap flow measurements

Sap velocity (vsap) was monitored half-hourly from 1 January 2013 to 31 October 2015 in eleven trees: six *E. obliqua* (mean diameter at breast height, DBH = 41 cm, range 30.8 - 50.9 cm), and five *E. rubida* (mean DBH = 33.3 cm, range 22.8 - 46.7 cm). All trees were healthy and all canopies had access to direct sunlight. Crown class as identified by canopy position (Smith et al., 1997) was evenly distributed between species, with two intermediate and three sub-dominant trees per species, and one dominant *E. obliqua* (no dominant *E. rubida* were present). We utilized the heat pulse compensation method to monitor sap velocity (The HeatPulser, Edwards Industries, Taupo, NZ), and distributed four probes per tree with increasing implantation depth to cover the sap velocity gradient (mean sapwood depth: *E. rubida* 1.95 ± 0.42 cm; *E. obliqua* 1.92 ± 0.38 cm). Sap velocity was corrected for deviations from exact parallel spacing of the heater and the thermistor elements, and wounding size was determined for all probes at the end of the study period. In addition, tree cores were collected next to each probe to correct measured vsap for the individual gas and water fractions of the sapwood of each instrumented tree (Edwards and Warwick 1984). Sap velocities were calculated for each probe and then averaged per tree. Each probe was analyzed for velocity drifts relative to the other three probes per tree, and affected probes were excluded from the analysis (2 probes in total). Data gaps through intermittent probe failures were filled with a hierarchical system to avoid an offset in the signal if either the fastest or slowest sensors were not working: 1. Gaps of individual probes were filled based on the highest fit of a regression between the probe that needed filling and the other three probes in the same tree (a probe was only chosen when R^2^ > 0.5). If the fit with any probe of the same tree was below the threshold, then the best fit with a probe from all other trees was chosen to fill the gap. 2. The vsap means of each tree were gap-filled if there were periods with a data gap affecting all probes of a tree simultaneously (e.g. during a power outage, during data downloads or sensor repairs). Here, we correlated the means of all trees with each other and chose the tree with the best fit. The lowest correlation between two trees had a Pearson R of 0.85, so no minimum threshold had to be applied.

### 2.3 Eddy covariance measurements

We monitored ecosystem-scale carbon and water exchange from a flux tower adjacent to the trees that were monitored for vsap dynamics. Fluxes were recorded at 30 m height with an open-path CO2/H2O analyzer (LI-7500, LI-COR Biosciences, Lincoln, NE, USA) and a 3-D sonic anemometer (CSAT3, Campbell Scientific Inc., Logan, UT, USA), sampled at 10 Hz and averaged over 30 minute intervals. Flux data were processed with ‘OzFluxQC’ version 2.9.5 (https://github.com/OzFlux/OzFluxQC), which included outlier removal through de-spiking, 2D co-ordinate rotations, WPL correction (Webb et al., 1980), conversion of virtual to sensible heat flux, and linear corrections for calibration anomalies and sensor drift (Griebel et al., 2017). We used the built-in neural network from OzFluxQC (SOLO; Isaac et al., 2017) for gap-filling of meteorological variables and fluxes. Data gaps of up to three hours were filled using linear interpolations, while longer gaps were filled with a descending preference of using alternative weather station data (Ballarat Aerodrome, ca. 20 km from study site), ACCESS model output from the Bureau of Meteorology, and BIOS2 model output, and lastly, using site-specific half-hourly averages of monthly climatology data (Isaac et al., 2017). We used 90-day intervals to gap-fill drivers and monthly intervals to gap-fill fluxes. Fluxes of carbon, water, and latent and sensible heat were gap-filled using incoming shortwave radiation, specific humidity deficit, and soil temperature as independent variables. Year-specific friction velocity thresholds were determined with the change-point detection method using 1000 iterations following Barr et al. (2013), which were subsequently applied for partitioning of net ecosystem exchange (NEE) into gross primary productivity (GPP) and ecosystem respiration (ER). Here, the neural network was trained with soil temperature and soil moisture as independent variables, before ER was predicted across a range of conditions that cover the entire data set of flux tower measurements. GPP was derived as GPP = – NEE+ER, where −NEE = NEP (net ecosystem productivity). We converted the gap-filled latent heat flux (Wm^−2^) to evapotranspiration (ET; mm) and daily means of water use efficiency (WUE) were derived as WUEday = ∑ GPP / ∑ ET (g C kg H2O^−1^; Table 1). Further, we calculated the daily mean underlying WUEu_day = GPPxVPD^0.5^/ET (gC kPa^−0.5^ kgH2O^−1^) to account for the non-liner effect of increasing VPD on LE (Zhou et al., 2015, 2014) and the daily mean intrinsic WUEi_day = GPP/Gc (gC mol^−1^ m^−2^ s^−1^) to quantify photosynthesis in relation to conductance (Beer et al., 2009; Lloyd et al., 2002; Schulze and Hall, 1982).

We calculated canopy conductance (Gc; mol m^−2^ s^−1^) and potential evapotranspiration (PET; mm) following Monteith (1965), where we used site-specific meteorological observations from the flux tower measurements as input parameters, and identified the well-watered reference surface conductance as the average of the surface conductance when VPD was between 0.9 and 1.1 kPa and surface soil moisture exceeded the 75% quantile. To account for the energy imbalance when inverting the Penman-Monteith equation to calculate canopy conductance and PET (Knauer et al., 2018; Wilson et al., 2002), we set the available energy equal to the sum of the latent and sensible heat flux, which implicitly conserves the Bowen ratio (Wohlfahrt et al., 2009). Rainy periods were excluded from the analysis, and half-hourly data were resampled to hourly periods to reduce noise.

### 2.4 Climate variables and response functions

In addition to fluxes, we measured the following meteorological variables above the canopy: downwelling and upwelling, shortwave and thermal radiation (CNR1; Kipp and Zonen, Delft, The Netherlands), air temperature and relative humidity (Vaisala HMP155; Vaisala, Helsinki, Finland). Precipitation was recorded as half-hourly totals at 1 m below the canopy (TB6; Hydrological Services Pty Ltd, Warwick Farm, Australia), and we added an additional rain gauge of the same type above the canopy in July 2014. Soil moisture measurements were initially recorded only at 10 and 40 cm depth close to the flux tower using time-domain measurement method to calculate soil volumetric water content (CS-616; Campbell Scientific Inc., Logan, UT, USA), and we extended these measurements by three additional sites adjacent to instrumented trees in November 2013. Thereafter, each of the four sites contained a CS-650 at 10 cm depth (Campbell Scientific Inc., Logan, UT, USA) and three additional CS-616 at 40 cm, 65 cm and ca. 1 m depth (depending on soil texture), and measurements from the four pits were averaged for each depth. We used one-way ANOVAs followed by a Tukey test to assess significant differences in the response and key climate variables between the baseline, the driest and the hottest days and heatwave days. All data were analyzed in R version 3.5.1 (R Core Team, 2018) using the packages ‘dplyr’ and ‘reshape2’ for manipulations, and ‘car’ for statistical analyses.

## 3 Results

### 3.1 Trade-offs between water loss and carbon gain

The daily sums of vsap, ET, GPP and daily means of WUE differed significantly among the baseline days, the hottest and the driest days, and during the heatwave (P < 0.01; Table 1). However, no variable was significantly different between the hottest days and the heatwave (Table 1), suggesting that the eucalypts did not respond differently to the longer heatwave compared with the individual hot days. On the driest days, vsap of both species remained comparable to the baseline, whereas vsap increased by >45% during the hottest days and by >70% during the heatwave (Table 1). Daily ET patterns resembled vsap patterns, indicating that transpiration dominated the ET signal of this forest. Daily GPP was comparable between baseline days, the hottest days and during the heatwave, but was significantly reduced during the driest days (16% lower than the baseline days; Table 1). However, daily ER relative to the baseline increased by 36% during the driest days, doubled during the hottest days, and increased by 137% during the heatwave (resulting in significant differences among all groups with the exception of the hottest and heatwave days; Table 1). This led to significant reductions in daily NEE relative to the baseline, which were in the order of 62% on the driest days, and 91% on the hottest days, and turned the forest from a moderate carbon sink to a carbon source (positive NEE) during the heatwave (Table 1).

Mean daytime canopy conductance relative to the baseline decreased significantly when soil moisture decreased (44% decrease in Gc during driest days with comparable mean daytime VPD; Table 1), and decreased further when VPD and temperatures increased during the hottest days and the heatwave (71% decrease in Gc despite comparable soil moisture to the baseline, P < 0.01; Table 1). WUE estimations were most sensitive to the method of calculation on the hottest days and during the heatwave (up to 91% difference within the same group; Table 1). Using the total WUEday, the significant difference in GPP did not translate to significantly different WUEday between the baseline and the driest days, which remained at 3 g C kg H2O^−1^ due to a comparable decrease in ET during the driest days. In contrast, WUEday decreased by 34% (2.0 ± 0.01 g C kg H2O^−1^) on the hottest days and by 42% (1.77 ± 0.05 g C kg H2O^−1^) during the heatwave, which resulted in significantly lower WUE during hot days than on the driest and baseline days. However, the opposite trend occurred when accounting for the non-linear relationship between GPPxVPD and ET at the ecosystem scale; that is, the underlying WUEu_day increased by 24% during the driest days (P<0.01; Table 1). Increasing WUE relative to baseline days was even more evident when based on the intrinsic WUEi_day, which increased by 30% during the driest days and by up to 125% during the hottest days and heatwave (P<0.01; Table 1), indicating that carbon assimilation (approximated as GPP) per unit stomatal conductance (approximated as Gc) significantly increased during high temperatures and high VPD, a result that was not captured when using total WUEday.

### 3.2 The diurnal cycle in response to increasing VPD

To further assess photosynthesis and water use during summer, we compared diurnal courses of Gc, GPP, vsap and ET as a function of VPD (Fig. 2a-e). On the hottest days, maximum vsap increased significantly (by 22% for *E. obliqua* and 28% for *E. rubida;* P<0.01) compared with baseline days, and vsap remained at these elevated levels while VPD was between 2.6 and 3.2 kPa. Thereafter, as VPD increased to 3.7 kPa in the mid-afternoon, vsap decreased by 20% for *E. obliqua* and 16% for *E. rubida* (i.e. equal to the maximum vsap on baseline days). vsap rates peaked during the heatwave (39 cm h^−1^ for *E. obliqua* and 47.4 cm h^−1^ for *E. rubida*) and remained at these significantly elevated levels (34% for *E. obliqua* and 42% for *E. rubida* above their baseline maximum; P<0.01) until VPD exceeded 4.3 kPa. In contrast, on the driest days, increasing VPD decreased the peak vsap of *E. obliqua* by 5%, while the peak vsap for *E. rubida* remained comparable to the baseline maximum (Fig. 2a, b). Apart from this small deviation on the driest days, both species had a similar response to changes in VPD.

**Figure 2.**
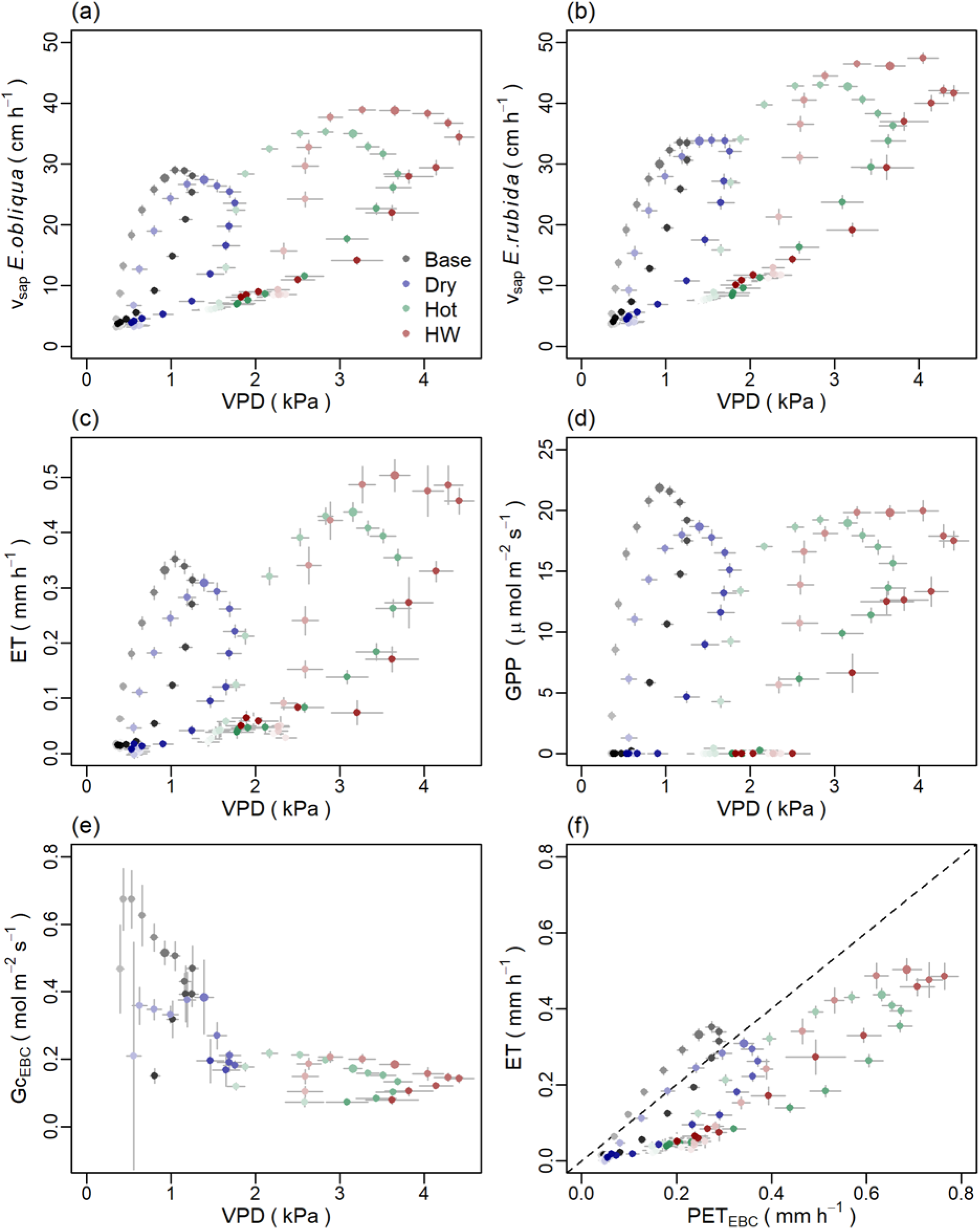
Diurnal patterns (means ± standard error) of sap velocity (vsap, cm h^−1^), evapotranspiration (ET, mm h^−1^), gross primary productivity (GPP, µ mol m^−2^ s^−1^), and daytime canopy conductance (GcEBC, mol m^−2^ s^−1^) in response to increasing vapor pressure deficit (VPD, kPa; panel a-e) and of ET against potential evapotranspiration (PETEBC, mm h^−1^; panel f) during the heatwave (‘HW’, red circles; 13-17 January 2014) on the hottest days (‘Hot’, green circles; air temperature > 30.7 °C), driest days (‘Dry’, blue circles; SWC < 0.1 m^3^ m^−3^) and baseline days (‘Base’, gray circles; January 2013, 2014 and 2015, excluding a hot period in January 2013, the January 2014 heatwave, and the hottest and driest days). Note that the subscript ‘EBC’ indicates that we set the available energy equal to the sum of the latent and sensible heat flux to account for the energy imbalance when inverting the Penman-Monteith combination equation to calculate canopy conductance and potential evapotranspiration. Symbols are colored according to the time of day (hourly time steps, shades darken with progression of the day) and the highlighted symbol indicates noon.

Consistent with vsap dynamics, maximum ET at the ecosystem scale increased 24% on the hottest days and by 43% during the heatwave (Fig. 2c; P<0.05 for the heatwave). However, despite only minor or no reductions in maximum vsap on the driest days, maximum ET decreased by 12% on the driest days relative to the summer baseline (Fig. 2c). As VPD increased in the afternoon, ET decreased between 9 to 28% until maximum VPD was reached during all conditions. Overall, ET dynamics were similar to vsap dynamics, but more closely resembled the dynamics of *E. obliqua* than *E. rubida* due the decrease of ET with increasing VPD during the driest days.

In contrast to significant increases in peak vsap and ET during the hottest days and the heatwave, the diurnal maximum of GPP declined with increasing VPD (Fig. 2d) and concurrently decreasing Gc (Fig. 2e) under both hot and dry conditions relative to the summer baseline (by 12% and 15% on the hottest and driest days, P<0.01; and by 9% during the heatwave, P>0.05; Fig. 2d). Until maximum VPD was reached in the mid-afternoon, GPP decreased between 12 to 19% during all conditions (Fig. 2d). While reductions of peak GPP were comparable to reductions of peak ET during driest days (both decreased by 12%), the diurnal course of GPP and ET reversed during the heatwave: mid-day GPP was sustained within 9% of baseline days (P>0.05) at the cost of significantly increased peak ET (43% increase compared to the summer baseline, P<0.05).

Daytime canopy conductance varied considerably between the hottest, driest and baseline days (Fig. 2e): maximum Gc at baseline days was 0.67 mol m^−2^ s^−1^, which decreased by 43% and 68% on the driest and hottest days (to 0.38 and 0.22 mol m^−2^ s^−1^, respectively). Maximum Gc was comparable between hot days and the heatwave, despite a larger VPD range during the heatwave (Fig. 2e). However, Gc did not fully decline under any conditions, and the large reductions of Gc on the driest, and especially on the hottest days and during the heatwave (Fig. 2e) only marginally affected GPP (Fig. 2d). Consequently, reductions in Gc primarily restricted excessive water loss during warm days with high atmospheric demand, which is supported by the ∼50% reduction of ET compared to PET (Fig. 2f and Table 1) during the hottest days and during the heatwave. In addition, the close resemblance of diurnal ET and PET dynamics on the baseline and driest days indicate that the forest was only marginally water limited on these days (Fig. 2f).

### 3.3 Carbon and water fluxes during a heat wave

The mean conditions during the five-day 2014 heatwave (13 to 17 January) compared with the local baseline for January 2013, 2014 and 2015 were characterized by similar soil moisture content (0.13-0.21 m^3^ m^−3^ during the baseline and 0.13-0.19 m^3^ m^−3^ during the heatwave), but by 7% higher incoming radiation peak, markedly warmer minimum (11.5 °C above baseline) and maximum temperatures (14.7 °C above baseline), and ∼four-fold higher atmospheric dryness (VPD), which peaked at ∼4.6 kPa in the early afternoon during the heatwave (Fig. 3a-b and Table 1). These increases in temperature and VPD resulted in a 37% increase in peak vsap rates and a 70% increase in total daily water use compared to baseline days (averaged across both species; Fig. 3c and Table 1). While peak PET increased three-fold during the heatwave, peak ET only increased by 43% compared to baseline days (Fig. 3c), leading to a 74% increase in total daily ET (from 2.88 mm on baseline days to 5.02 mm during the heat wave; Table 1). In contrast to increases in vsap and ET, the daily peaks and daily totals of GPP remained relatively unchanged during the heatwave, indicating that baseline photosynthesis was maintained during the heatwave at the cost of significantly increased transpiration (Table 1 and Fig. 3d). However, despite stable GPP, a doubling of peak respiration rates (P<0.05) and a more than two-fold increase in daily ER (P<0.01) turned the forest from a moderate net carbon sink during baseline days (−4.64 ± 0.34 g C d^−1^) into a net carbon source (0.79 ± 0.47 g C d^−1^) during the heatwave (P<0.01; Fig. 3d and Table 1).

**Figure 3.**
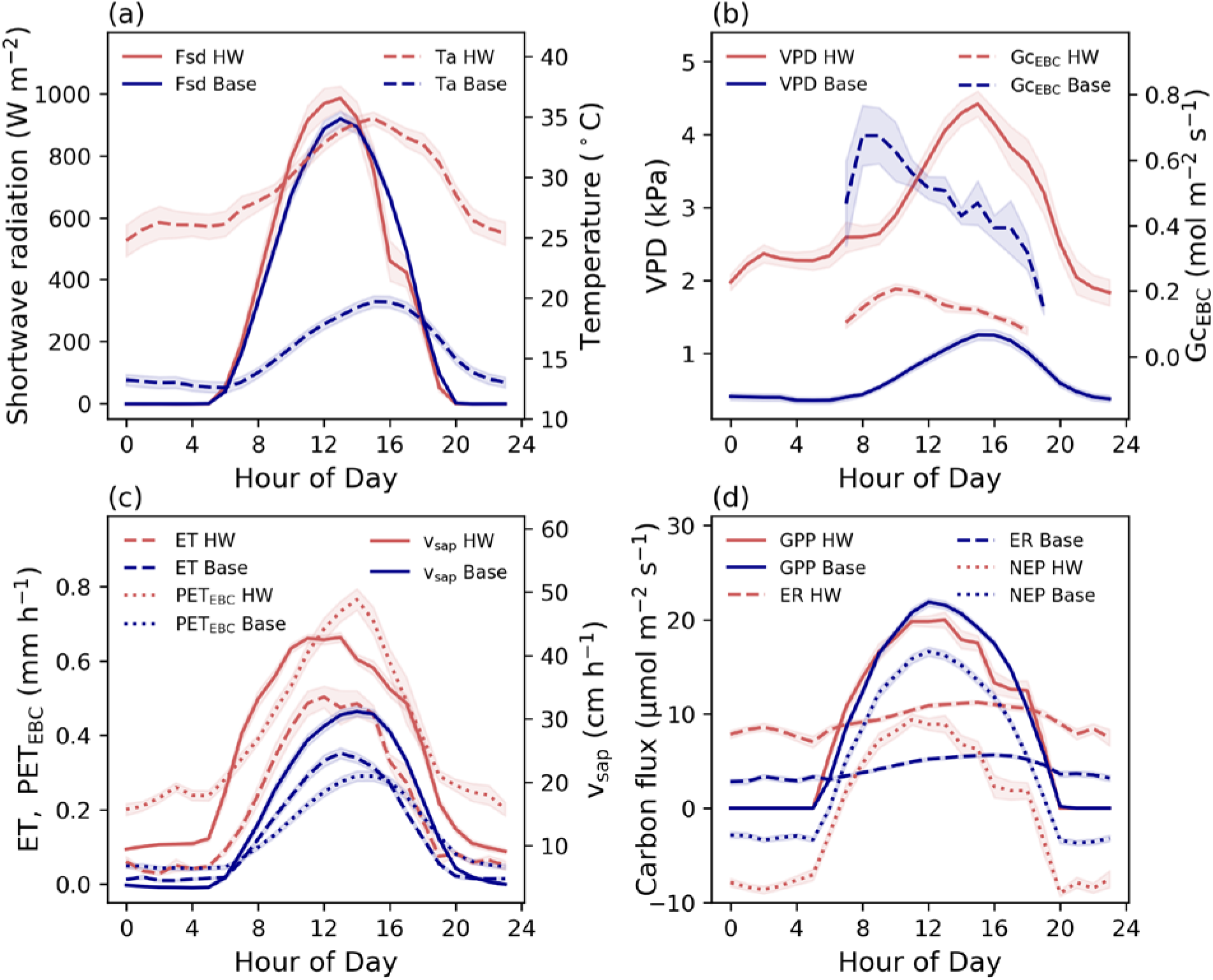
The mean conditions during the heatwave (HW, 13-17 January 2014; red lines) compared with the local baseline (Base, mean of January 2013-2015 without a hot period in 2013, the 2014 heatwave and the hottest and driest days; blue lines). Abbreviations: Fsd = shortwave radiation (W m^−2^), Ta = air temperature (°C), VPD = vapor pressure deficit (kPa), GcEBC_day = mean daytime canopy conductance (mol m^−2^ s^−1^), vsap = sap velocity (cm h^−1^, average of *E. obliqua* and *E. rubida*), ET = evapotranspiration (mm h^−1^), PETEBC = potential evapotranspiration (mm h^−1^), GPP = gross primary productivity (µ mol m^−2^ s^−1^), NEP = net ecosystem productivity (µ mol m^−2^ s^− 1^), ER = ecosystem respiration (µ mol m^−2^ s^−1^). Note that shading represents standard error, and the subscript ‘EBC’ indicates that we set the available energy equal to the sum of the latent and sensible heat flux to account for the energy imbalance when calculating PET.

## 4 Discussion

### 4.1 Increased water use to sustain photosynthesis during high temperatures

Contrary to our first hypothesis, photosynthesis and water use within this dry-sclerophyll eucalypt forest were not always synchronous during summer, as we measured no significant change in photosynthesis in contrast to significantly increased water use in daily sums (Table 1), in response to increasing VPD (Fig. 2), and in diurnal patterns (Fig. 3) during the hottest days and during the 5-day heatwave. On the hottest days and during the heatwave, photosynthesis was more-or-less sustained at the cost of >70% increased water loss relative to baseline days, which contrasted with concurrent decreases in both GPP (16%) and ET (16%) on the driest days. While hot or dry conditions resulted in decreased canopy conductance, this primarily restricted excessive water loss due to high atmospheric demand, and only marginally affected the GPP of this temperate eucalypt forest.

The ability to maintain photosynthesis across hot or dry conditions indicates a yield-focused growth strategy of the local eucalypt species, where carbon gain through photosynthesis is prioritized over a conservative water use; this could partly explain the high annual carbon sequestration rates that have been reported for this forest in comparison with other temperate eucalypt forests (Beringer et al., 2016; Griebel et al., 2017; Hinko-Najera et al., 2017). Moreover, our findings demonstrate potential for temperate eucalypt forests growing in relatively mild, mesic conditions similar to our study site (e.g. at elevation or on cold sheltered sites with sufficient moisture supply) to maintain photosynthesis under future warmer climates, whereas eucalypt forests in less favorable growing conditions will likely increase stomatal adjustment to levels that adversely affect photosynthetic uptake (Drake et al., 2018; van Gorsel et al., 2016; Renchon et al., 2018). However, a doubling of ecosystem respiration turned the forest from a moderate net carbon sink into a net carbon source during the heatwave. Thus, while the photosynthesis of this eucalypt forest appears largely unaffected during warmer and drier conditions in the summer months, the over-proportional increase in ER resulted in a switch from a net sink to a source, highlighting that the net productivity can be adversely affected by isolated extreme events. Hence, with a projected increase in the number, duration and intensity of heat waves, the potential of temperate eucalypt forests to remain carbon sinks under future climates will largely depend on the response of ecosystem respiration to increasing temperatures rather than the ability to sustain photosynthesis during extreme heat.

The large increase in ET during the hottest days and the heatwave resulted in up to two-fold variations in total WUE in summer, with conflicting trends depending on the formulation of WUE (Table 1). While seasonal variation in WUE within the same ecosystem is typically linked to variations in canopy phenology (Huang et al., 2016; Jin et al., 2017), our observations were constrained to the summer season and pooled across three different summers with similar radiation input for all examined conditions (Fig. S1.), thereby minimizing phenological influences as well as climatological variation. Yet, significant differences between the baseline and driest days in underlying WUE indicated that this metric was more sensitive to capturing physiological responses to drought stress than intrinsic WUE, possibly because WUEu_day reduces large parts of the diurnal and seasonal variation at the ecosystem scale within the same plant functional type (Zhou et al., 2015). In contrast, WUEi_day doubled during the hottest days and the heatwave, and accurately captured the significantly increased rate of photosynthesis per unit conductance. Despite different sensitivities of underlying and intrinsic WUE to either dry or hot conditions at our study site, none of these physiological responses were captured using the traditional formulation of WUE (GPP/ET), supporting that alternative formulations of WUE improved insights into the mechanisms regulating water loss and carbon uptake.

### 4.2 Dependence on atmospheric demand

The influence of VPD on carbon and water fluxes has been well acknowledged (Beer et al., 2009; Eamus et al., 2013; Knauer et al., 2015; Novick et al., 2016; Renchon et al., 2018), and short- and long-term reductions in GPP and transpiration due to fluctuations in VPD and soil moisture have been recorded across a range of biomes even during non-drought years (Sulman et al., 2016). Further, a decrease in transpiration and photosynthesis due to stomatal regulation in response to increasing VPD is well established for eucalypts (Duursma et al., 2014; Mitchell et al., 2012; Pepper et al., 2008; Prior et al., 1997). While our study indicated that stomatal regulation of water loss can differ between exceptionally hot conditions and dry conditions, ecosystem-scale estimates of canopy conductance to infer stomatal activity are subject to large uncertainties as Gc is only inferred and not directly measured (Knauer et al., 2015; Wohlfahrt et al., 2009). While we attempted to minimize the contribution of non-transpirational water fluxes by removing rainy periods and by forcing energy balance closure (section 2.3), stomatal regulation might still be confounded by anatomical properties of water transport; that is, the vsap to VPD response can vary if the water absorption capacity at the root-soil interface becomes limiting or if the xylem anatomy restricts the water transport capacity from the roots to the canopy (Eamus & Prior, 2001; Sperry & Pockman, 1993; Tyree & Ewers, 1991). Still, ET was significantly reduced compared with PET on the hottest days and during the heatwave, providing clear indications of physiological regulations of water loss in response to increasing atmospheric demand in our forest, whereas the relative importance of stomatal regulations versus hydraulic restrictions on water transport remains unclear. Furthermore, the ability to maintain or even increase vsap rates in all the conditions of our study suggests that the trees likely had access to soil water in deeper layers (section 4.3). Despite excluding rainy periods from our analyses, reduced peak rates and daily totals of ET on the driest days were not clearly due to decreases in vsap, indicating a greater relative contribution of soil and canopy evaporation (or the lack thereof) to ecosystem-scale ET dynamics during dry conditions. Nevertheless, ET dynamics closely resembled vsap dynamics of *E. obliqua*, confirming that stand-scale observations of ET were dominated by the transpiration dynamics of the dominating species (70% of stand basal area; Griebel 2016).

### 4.3 Dependence on water availability

We can partially confirm our second hypothesis that evapotranspiration increases during a longer heatwave, however GPP remained comparable to baseline days and we did not measure a simultaneous sharp reduction in photosynthesis rates during the heatwave (as was measured for one-year old eucalypt saplings by Drake et al., 2018). This suggests that temperature stress may be ameliorated by water access at this comparatively mesic site. In addition to moderate moisture levels down to 1 m depth during the heatwave (ranging from 0.13 to 0.19 m^3^ m^−3^), sustained vsap even during the driest days suggests that the trees had access to deep water reserves, which were likely recharged in the 2010/2011 LaNiña years when annual rainfall totals were ∼200-400 mm above the long-term average. Eucalypts can have fast-growing roots that allow them to reach deep water resources quickly (e.g. 12 m depth in 2 years; Christina et al., 2017), and deep soil water access is known to be an important buffer in north Australia’s open forests and savanna regions, where large amounts of water stored during the wet season can be accessed by trees during dry periods (Arndt et al., 2015; Eamus et al., 2015; Hutley et al., 2000; O’Grady et al., 1999). Even for temperate eucalypt forests in more complex terrain, access to water in deep soil layers has explained water losses that have exceeded annual precipitation inputs (Benyon & Doody, 2015; Mitchell et al., 2012). Hence, while access to deep soil water might represent an efficient adaptation of eucalypt trees to drought (Duursma et al., 2011; Markewitz et al., 2010; Nepstad et al., 2007; Yang et al., 2017), our study indicates that deep soil water access also plays an important role in explaining comparatively high carbon sequestration rates of temperate eucalypt forests. Furthermore, we found no evidence that the forest was exposed to severe heat or drought stress during our three observation years, which is supported by an absence of increased leaf shedding (Griebel et al., 2015) - a typical stress response for eucalypts (Granda et al., 2014; Pook, 1984; Renchon et al., 2018; Silva et al., 2004) - and by a sustained leaf area index whereby leaf loss was consistently balanced by growth of new leaves in our forest (Griebel et al., 2017). In addition, sustained photosynthetic rates during high temperatures indicate some buffer in the capacity of this forest type to maintain productivity under a warming climate; however, effects on productivity of very hot conditions combined with very dry conditions remain unclear, particularly if sustained dry conditions lead to a depletion in deep soil water reserves.

## 5 Conclusions

We present evidence that this temperate eucalypt forest was able to sustain photosynthesis at the cost of increased water loss during individual exceptionally hot days and during a longer heatwave, which contradicted our hypotheses of (i) concomitant reductions or increases in transpiration and photosynthesis, and (ii) notably decreasing photosynthesis with increasing transpiration during heatwaves. Increased or sustained transpiration rates during the hottest, driest and heatwave days indicated sufficient water availability to sustain photosynthesis rates at our study site, and consequently that neither individual hot or dry days, nor the heat wave coincided with water deficit and/or drought stress. This, in turn, indicated access of the roots to deeper water reserves. How sustainable such water reserves are, and how much these reserves will be depleted by prolonged heat waves or dry conditions remains unclear. Moreover, the switch from a net sink to a net source of carbon during the heatwave highlights potential limitations to similar, high elevation temperate eucalypt forests remaining carbon sinks under future climates, which will largely depend on the response of ecosystem respiration to increasing temperatures rather than their potential to maintain photosynthesis during hot or dry conditions.

## Acknowledgments

We thank John Collopy and Julio Najera-Umana for assisting with the installation of the field equipment. This study was partly funded by TERN-OzFlux and TERN-SuperSites, the Australian Research Council (ARC) grants LE0882936 and DP120101735, and the Integrated Forest Ecosystem Research program supported by the Victorian Department of Environment Land, Water and Planning (DELWP). The flux tower data are available at the OzFlux portal (http://data.ozflux.org.au) and the tree water use data are available at Sapfluxnet (http://sapfluxnet.creaf.cat/app). We appreciate substantive input from four anonymous reviewers which improved the quality of the manuscript.

## Supporting Information

**Figure S1.**
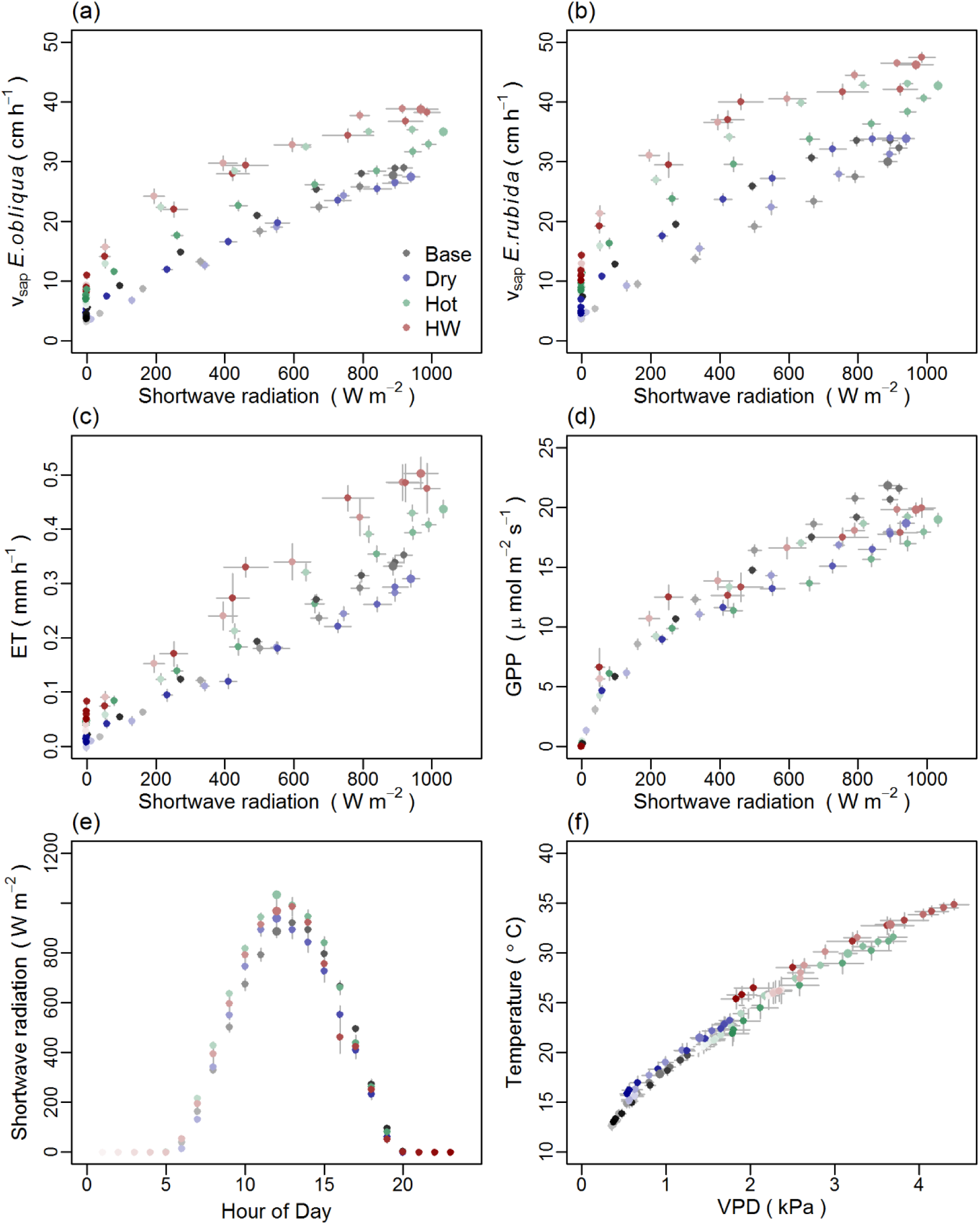
Diurnal patterns (means ± standard error) of sap velocity (vsap, cm h^−1^), evapotranspiration (ET, mm h^−1^), gross primary productivity (GPP, μ mol m^−2^ s^−1^) against solar radiation (Shortwave radiation, W m^−2^; panel a-d), as well as the diurnal course of shortwave radiation (panel e) and of VPD against air temperature (°C; panel f) during the heatwave (‘HW’, red circles; 13-17 January 2014) on the hottest days (‘Hot’, green circles; air temperature > 30.7 °C), driest days (‘Dry’, blue circles; SWC < 0.1 m^3^ m^−3^) and baseline days (‘Base’, gray circles; January 2013, 2014 and 2015, excluding a hot period in January 2013, the January 2014 heatwave, and the hottest and driest days). Symbols are colored according to the time of day (hourly time steps, shades darken with progression of the day) and the highlighted symbol indicates noon.

